# DeepRBP: A novel deep neural network for inferring splicing regulation

**DOI:** 10.1101/2024.04.11.589004

**Authors:** Joseba Sancho, Juan A. Ferrer-Bonsoms, Danel Olaverri-Mendizabal, Fernando Carazo, Luis V. Valcárcel, Idoia Ochoa

## Abstract

**Motivation:** Alternative splicing plays a pivotal role in various biological processes. In the context of cancer, aberrant splicing patterns can lead to disease progression and treatment resistance. Understanding the regulatory mechanisms underlying alternative splicing is crucial for elucidating disease mechanisms and identifying potential therapeutic targets.

**Results:** We present DeepRBP, a deep learning (DL) based framework to identify potential RNA-binding proteins (RBP)-Gene regulation pairs for further in-vitro validation. DeepRBP is composed of a DL model that predicts transcript abundance given RBP and gene expression data coupled with an explainability module that computes informative RBP-Gene scores. We show that the proposed framework is able to identify known RBP-Gene regulations, demonstrating its applicability to identify new ones.

**Availability and Implementation:** DeepRBP is implemented in PyTorch, and all the code and material used in this work is available at https://github.com/ML4BM-Lab/DeepRBP.

**Supplementary information:** Supplementary data are available at *Bioinformatics* online.

## Introduction

Alternative splicing (AS) is a fundamental molecular process where exons, the coding regions of the genome, are diversely assembled to produce distinct isoform variants from a single gene (Oltean and Bates, 2014). This variation can lead to the generation of multiple different proteins from a single gene, greatly increasing the protein and functional diversity of an organism. This post-and co-transcriptional process is a critical research area, as aberrant splicing is related to tumor and tissue phenotypes (Oltean and Bates, 2014). Highly regulated and orchestrated, AS relies on specific RNA-binding proteins (RBPs), called splicing factors, that together with the spliceosome complex, remove introns and connect exons, ensuring the proper assembly of mRNA transcripts (Dominguez *et al*., 2018; Hong, 2017).

Unfortunately, experimental methods for characterizing RBPs and their splicing targets, such as CLIP experiments, are expensive and labor-intensive (Ramanathan *et al*., 2019). Moreover, it is important to note that while these methods can demonstrate the binding of an RBP to RNA, they may not conclusively prove whether the RBP has acted on splicing (Ramanathan *et al*., 2019).

Given the requirement of a targeted RBP for conducting a CLIP experiment, several algorithms have been developed to predict the potential involvement of specific RBPs in splicing interactions, as they could be used as a new therapeutic targets to treat diseases (Hong, 2017; Li *et al*., 2021; Carazo *et al*., 2019; Park *et al*., 2016; Chen and Keleş, 2020). These methods integrate RBP databases, such as POSTAR3 (Zhao *et al*., 2022), and a statistical analysis of AS performed by methods such as EventPointer (Ferrer-Bonsoms *et al*., 2022), rMATS (Shen *et al*., 2014) or SUPPA (Sebestyén *et al*., 2016). However, these algorithms are restricted by the number of RBPs available in the CLIP experiment databases. Furthermore, they do not take into account the possibility that splicing regulation may be orchestrated by the simultaneous interaction of multiple RBPs or even the possibility that one RBP regulates the activity of other RBPs (Gárate-Rascón *et al*., 2022).

Deep learning (DL) approaches have also emerged as a promising solution to study AS. For example, in (Zhang *et al*., 2019) they proposed a method based on DL to infer differential AS from bulk RNA-Seq data in two conditions. One of the main challenges in the field of DL is the explainability of the predictive models, often referred to as the “black box” problem. Although this remains a significant hurdle, methods such as

Deep Learning Important FeaTures (DeepLIFT) (Shrikumar *et al*., 2017), which predicts the importance of each input variable in the model’s prediction, can lead to a better understanding of how a DL model works.

To our knowledge, there are no approaches that predict splicing regulation from bulk RNA-Seq data. In this work, we propose DeepRBP, a DL-based framework that identifies potential regulations in the form RBP-Gene (or RBP-Transcript) given bulk RNA-Seq data. DeepRBP is composed of a trained DL model that predicts transcript abundance given RBP and gene expression data, coupled with an explainability module based on DeepLIFT that generates scores for every RBP-Transcript pair. These scores are then used to identify potential RBP-Gene regulations. DeepRBP therefore leverages an interpretable model with the aim of unveiling novel regulatory relationships for subsequent in-vitro validation. We show through extensive simulations that the proposed model is able to correctly infer transcript abundance, and more importantly, to identify known RBP-Gene regulations. Moreover, we use the proposed scheme to uncover potential unknown RBP-Gene regulations in acute myeloid leukemia (AML), kidney chromophobe (KICH), and hepatocellular carcinoma (HCC).

## Materials and Methods

### Considered RBPs and Genes

The set of considered genes was compiled by merging three distinct data-sources: genes whose alternative splicing has been demonstrated to contribute to cancer (*N* = 89) (Table S5, Sebestyén *et al*. (2016)), genes predicted to be potential cancer drivers based on mutations and/or copy number alterations (*N* = 900) (Table S6, Sebestyén *et al*. (2016)), and genes with published evidence in oncology from the Catalogue of Somatic Mutations in Cancer (COSMIC) (*N* = 736) (Sondka *et al*., 2018). Additionally, only genes comprising more than one transcript were considered, as it is necessary for alternative splicing. Altogether, this resulted in 11,462 transcripts and 1,040 genes (see Supplementary Fig. 1 and Supplementary table 1.xlsx).

The list of considered RBPs was taken from Sebestyén *et al*. (2016) (Table S2), containing a total of 1,282 reviewed RBPs (Supplementary table 2.xlsx).

### DeepRBP Prediction Model

#### Model Architecture

The prediction module aims to predict transcript abundances using the expression of the RBPs. To achieve this, the RBP expression data is passed through fully-connected layers followed by batch normalization layers, as described in (Ioffe and Szegedy, 2015), and activation layers. The final architecture is selected via an optimization process (see Model Optimization below). At the output layer, the sigmoid activation function is applied to obtain the percentage of the gene expression that contributes to each transcript, such as the predicted percentages for all transcripts of a gene should sum up to one.

This formulation, however, may not be representative for low-expressed genes, potentially harming the training of the model, which is performed by minimizing the Mean Squared Error (MSE) between the predicted and real values. For this reason, we introduce the expression of the genes as input, and compute instead transcript abundances, i.e., the transcripts’ expression. In particular, the predicted percentages are multiplied by the corresponding gene expression, input in TPMs, yielding transcript abundances also in TPMs. Note that this transformation is applied solely to the computation of the cost function (MSE in this case), ensuring that low-expressed genes have a low influence in the overall error (free percentages). Additionally, we apply the log_2_(*x* + 1) transformation so that moderately and highly expressed genes contribute equally to the error. In summary, the loss is computed as

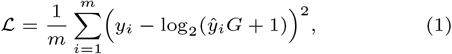

where *m* is the number of transcripts, *y*_*i*_ is the measured transcript expression in log_2_(*TPM* + 1), *ŷ*_*i*_ is the percentage predicted by the output layer, and *G* is the expression of the associated gene (in TPM).

#### Model Optimization

Hyperparameter tuning and architecture optimization were carried out by experimenting with the hyperparameters described in Table 1. Note that 0 hidden layers reduces to logistic regression models to predict transcript percentages. The activation function is applied to all hidden layers, and *node number scaler* reduces the number of nodes from layer to layer by that amount if *number nodes constant* is true, otherwise, the number of nodes is reduced only in the last hidden layer (if more than 1 hidden layer). The number of nodes in the last hidden layer is reduced to be consistent with the biological principle that RBPs tend to aggregate in small groups to regulate the splicing process (Carazo *et al*., 2019). Note that the size of the last hidden layer can be at most 2, 048, much smaller than the number of considered transcripts (11, 462).

**Table 1.** Summary of the considered hyperparameters for architecture optimization.

| Hyperparameter | Tried values |
| --- | --- |
| Activation function | ReLU, tanh, or sigmoid |
| Number of hidden layers | 0, 1, 2, 3 |
| Number of nodes in first layer | 128, 256, 512, 1024, 2048 |
| Node number scaler | 2, 4, 8 |
| Number of nodes constant | True, False |
| Learning rate | 0.0001, 0.001, 0.01 |

Optimization was performed using Optuna (Akiba *et al*., 2019), a tool to accelerate the hyperparameter tuning process by suggesting values within predetermined ranges. Its built-in function TPESampler (Bergstra *et al*., 2011, 2013) was applied to select the next hyperparameters to be assessed. In total, we performed 100 iterations with TPESampler, thus training 100 different models. AdamW optimizer was used for training with varying learning rate (see Table 1), a batch size of 128, 1,000 epochs, and MSE as loss (see Model Architecture above). This training configuration is also used to train the final model with the best hyperparameters.

The selection criteria for determining the optimal combination of hyperparameters involved ranking the models based on the following key metrics on the validation set: Spearman’s coefficient and MSE. Additionally, models with a lower parameter number in its architecture were given preference.

#### Final Prediction Model

To define the model architecture and train the final DeepRBP prediction model we considered data from The Cancer Genome Atlas (TCGA) (Weinstein *et al*., 2013), available from the the University of California Santa Cruz (UCSC) Xena platform (https://xenabrowser.net/) (Goldman *et al*., 2020). The TCGA database contains samples from 33 different cancer types spread in 9,185 samples with matched 727 normal samples. 80% of samples from each tumor type, with both tumor and match normal samples, were considered as the training set, while the remaining 20% were set aside for testing.

Model optimization was carried out using half of the TCGA training data, aiming to optimize time and computational resources. Within this subset, 80% (*n* = 2, 678) was allocated for training and the rest (*n* = 682) for validation (see Supplementary Table 1 for details on the number of samples per tumor type). This led to an architecture composed of two hidden layers with 1024 and 128 nodes, respectively, ReLu activation function, and a learning rate of 0.001 (see Results section for details). An schematic of the considered architecture is depicted in Fig. 1A. The TCGA training set was used to train the final model. As shown in the results section, training with samples from all tumor types increases the generalization ability of the model, facilitating the use of the proposed framework to infer unknown regulations, as there is no need to retrain the prediction model.

**Fig. 1.**
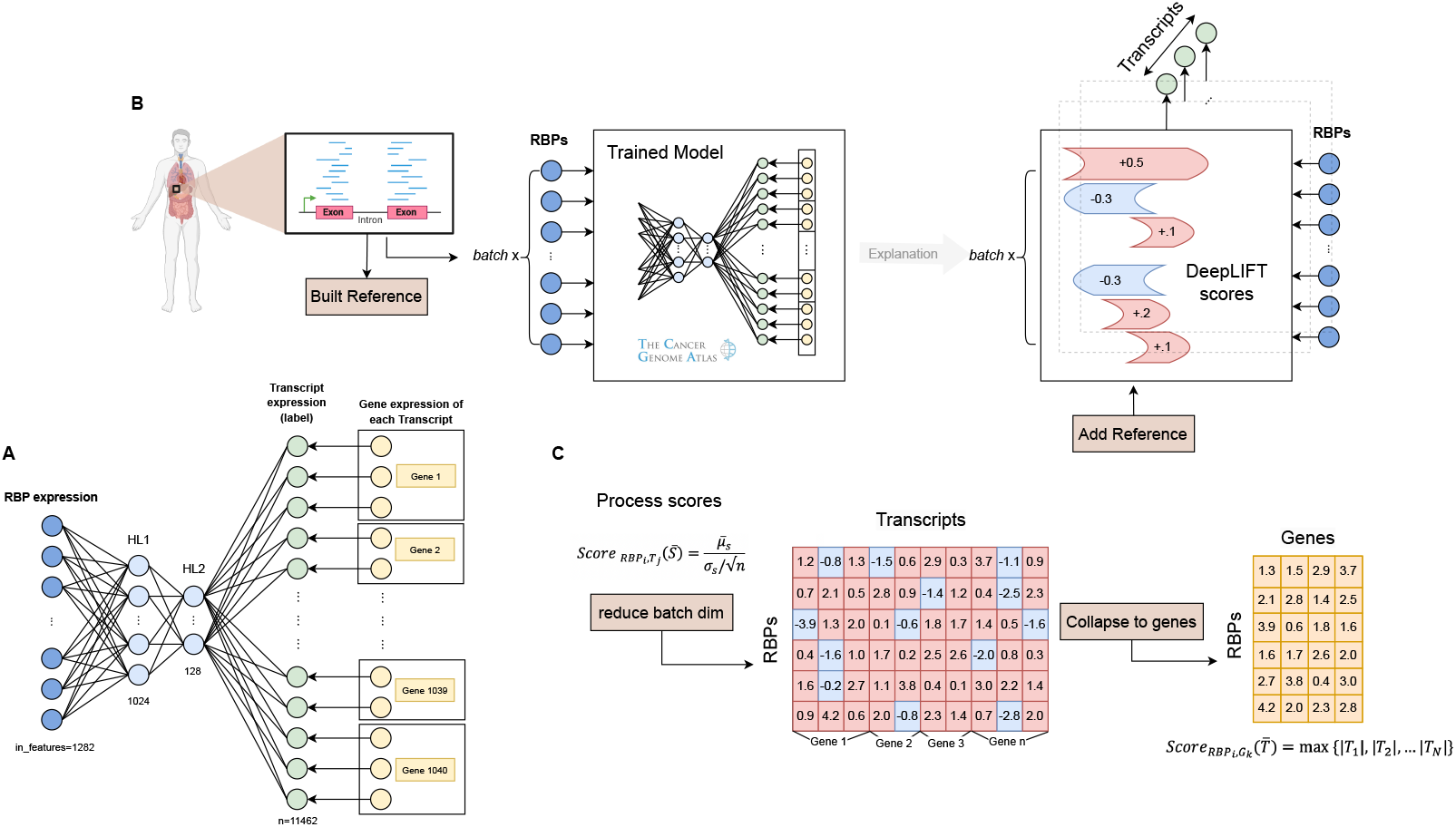
Overview of the DeepRBP framework. **A**. The DL-based predictive model is trained to learn the isoform usage (transcript abundance) from processed RBP and gene expression data as a proxy to identify potential unknown RBP-Gene regulations. The model is trained on TCGA data comprising several cancer types. **B**. Given RNA-Seq data of a specific cell line or tissue and a built reference, scores for each RBP-Transcript pair and sample are computed with DeepLIFT using the TCGA-trained model. **C**. Scores along the sample dimension are condensed using a z-score to obtain a score matrix of RBPs by transcripts. Subsequently, transcript-level scores are collapsed to genes by taking for each gene the maximum absolute score among its transcripts. Higher scores are associated with potential RBP-Gene regulations.

The model is implemented in PyTorch and publicly available at https://github.com/ML4BM-Lab/DeepRBP. While users have the option to train the model from scratch, the pre-trained model using TCGA data is provided within the DeepRBP framework and ready to use.

### DeepRBP Explainability Module

We used DeepLIFT (Shrikumar *et al*., 2017), a method for model explainability, to understand how input features (RBPs in our case) contribute to the outputs (transcripts in our case) of the final trained model. Specifically, given input RNA-Seq data, e.g., from a specific cell-line or tumor type, and a reference, DeepLIFT computes RBP-Transcript scores. We use a knockout reference (an all-zero vector) in our experiments, as using the median resulted in less-interpretable scores (see Results Section).

Using DeepLIFT, the contribution of each RBP-Transcript pair is determined for every sample in the input data, obtaining a three-dimensional score matrix with dimensions: number of transcripts by number of RBPs by number of samples (see Fig. 1B). Positive scores indicate activation of the transcript, while negative scores indicate transcript inhibition. To obtain a single score indicative of the general behavior of each RBP-Transcript pair, we collapse the scores across samples by computing the t-statistic given by 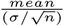, where *n* indicates the number of samples. This results in a score matrix of size number of transcripts by number of RBPs. Collapsing by summing the scores yielded less-informative scores (see Results section).

The RBP scores at the transcript level can further be collapsed to genes by taking, for each gene, the maximum absolute value across its transcripts, resulting in a gene by RBP score matrix (see Fig. 1C). We considered the absolute value as we are interested in identifying RBP-Gene regulations, independently of the sign. This transformation to the gene level implies, according to the maximum of a distribution of random variables, that genes with more transcripts will have a higher probability of obtaining higher RBP scores. This aligns with the biological intuition that a greater abundance of transcripts indicates a more substantial impact on the splicing process.

### Data Preprocessing

For all considered datasets in this study, the expression of the selected RBPs is transformed to log_2_(TPM + 1). Subsequently, the RBP data is normalized to z-scores per RBP, subtracting the mean and dividing by the standard deviation. In order to fall within the range of zero to one, z-scores are truncated to (−2, +2) and linearly displaced to the desired range. The rationale to truncate the values to (−2, +2) was to avoid outliers (see Supplementary Fig. 4).

The transcript expression of the considered genes is also transformed to a log_2_(TPM + 1) scale, to align with the loss computation of the prediction model (see model architecture), while the gene expression is input to the model in TPM.

### DeepRBP Validation

#### Generalization ability of the prediction model

To assess the generalization ability of our prediction model across various tumor types and cellular conditions, we utilized the 20% subset of TCGA samples from all tumor types set aside (*n* = 2121) alongside healthy samples from the Genotype-Tissue Expression (GTEx) (Lonsdale *et al*., 2013) database. Incorporating GTEx data is pivotal for evaluating the model’s performance on healthy samples, despite observing some batch effects compared to the TCGA samples (see Supplementary Fig. 5). The GTEx data was also downloaded from the UCSC Xena platform, and it contains gene expression profiles of 54 specific tissues and 3,988 samples.

#### Score validation with POSTAR3

To validate the scores obtained with DeepLIFT, we used human POSTAR3 (Zhao *et al*., 2022), a comprehensive Post-Transcriptional Regulation database. POSTAR3 provides protein binding sites on RNA obtained by CLIP experiments. Starting from the binding site data of the RBPs in the genome, the event (*E*) information, and the regions of these events, we match and create a binary *E × RBP* matrix. A value of 1 indicates that an event is regulated by the corresponding RBP, while a 0 indicates no regulation, and an empty (N/A) value corresponds to unknown regulations. From the *E ×RBP* matrix we derive the *G × RBP* matrix by grouping all events in a gene and limiting values to 0, 1, and N/A (*G* stands for genes). In other words, for a given gene and RBP, if at least one event of the gene is regulated by the RBP, we put a 1 in the *G × RBP* matrix at that position. We did not compute a transcript by RBP matrix since it is not straightforward to relate events to transcripts. In the following, we denote by *class 1* (*class 0* ) to all RBP-Gene pairs with a 1 (0) in a given *G × RBP* matrix.

Analyzing the POSTAR3 database, we observed that the tumoral cell-lines with most CLIP experiments and consequently more analyzed RBP-Events corresponded to HepG2, Huh7, K562 and HEK293, and hence we used data from these cell-lines to validate the proposed framework. We considered a single *G × RBP* matrix for HepG2 and Huh7, since both correspond to liver tumor (see Supplementary Table 2 and Supplementary Fig. 6 for a detailed description of the generated POSTAR3 binary matrices).

Three independent experiments with the proposed DeepRBP framework were then carried out. We utilized the following RNA-Seq samples from TCGA to compute the corresponding RBP-Gene scores: liver hepatocellular carcinoma (for HepG2 and Huh7), acute myeloid leukemia (for K562), and kidney chromophobe (for HEK293). For the last two cell lines, DepMap Celligner Warren *et al*. (2021) was employed to map the cell-line to the most similar TCGA tumor samples. Only primary tumor samples were used (*N* = 79, *N* = 35 and *N* = 19, respectively). Using the computed binary POSTAR3 matrix, we analyze whether the obtained scores with DeepRBP can identify the known RBP-Gene relationships (*class 1* ). We expect the class 1 RBP-Gene pairs to get higher scores than the *class 0* ones. We perform Mann-Whitney-Wilcoxon tests across RBPs or across genes. The p-value results were corrected using Bonferroni correction.

#### Score validation with real RBP knockdown data

We utilized DeepRBP to analyze RNA-seq data from PRJEB39343 (Cheng *et al*., 2020), GSE77702 (Kapeli *et al*., 2016), GSE75491 (Yang *et al*., 2016) and GSE136366 (Roczniak-Ferguson and Ferguson, 2019), which involved the knockdown of specific RBPs (see Table 2 for details). Briefly, PRJEB39343 experiments were conducted on gastric cancer cell lines, GSE75491 on the H358 cell line, GSE136366 on the HeLa cell line, and GSE77702 on human motor neurons derived from induced pluripotent stem cells.

**Table 2.** Details of real knockdown experiments.

| Project Name | knockdown RBP | Cell Line |
| --- | --- | --- |
| PRJEB39343 | MBNL1 | gastric cancer |
| GSE75491 | RBM47 | lung cancer |
| GSE77702 | FUS, TAF15, TARDBP | motor neurons |
| GSE136366 | TARDBP | cervical |

RNA-Seq data was processed with Kallisto, using Gencode23 as the reference transcriptome, to determine transcript expression. The gene expression was then calculated as the aggregate of the expression of its associated transcripts. The resulting counts were processed as previously described and then fitted into the trained DeepRBP model. The model’s performance in predicting transcript abundance was evaluated using Spearman’s correlation and MSE, for control and knockout samples separately.

For each experiment, we then identified which transcripts are affected by the knockout of the RBP, by performing differential expression (DE) analysis of control samples against knockout samples using voom-limma (Law *et al*., 2014) at the transcript level. We also identified the genes for which at least one transcript was differentially expressed. Subsequently, control samples were used with DeepRBP to obtain scores at the transcript and gene levels for each RBP. For each RBP, we then compared the scores between differentially expressed transcripts and those that were not through Mann-Whitney-Wilcoxon tests. We expect to see significant differences for the knocked RBP, but not for the other RBPs.

### Identification of potential RBP-Gene regulations

For the experiments related to the POSTAR3 database, we computed specific thresholds for each RBP based on the score distribution of the known 0s and 1s. The threshold is optimized to maximize the difference between the true positive rate and the false positive rate. Then, for unknown RBP-Gene regulations (specified as N/A in the POSTAR3 binary matrix), those that exhibit a score surpassing the RBP-threshold are identified as potentially regulated. These are therefore candidates for further in-vitro validation.

### Hardware

All simulations were performed on a cluster, using a server node with 48 cores Intel xeon gold 6248R with 96GB RAM and a NVIDIA RTX 3090 GPU (CUDA v11.2).

## Results

### DeepRBP model selection

The results obtained from performing the hyperparameter and architecture optimization are summarized in Supplementary Tables 3-4. Based on the obtained results, the hyperparameter configuration of the *#41* model was chosen for the final architecture of DeepRBP prediction model. This model ranks within the top 5 across both MSE and Spearman’s correlation metrics, obtaining a similar performance to the alternative configurations while maintaining a smaller parameter count. This is why, albeit model *#87* displaying the best results overall, we opted for model *#41*. We note also that the model with 0 hidden layers (baseline model) does not exhibit a high ranking. In fact, all top-ranked models have a minimum of two hidden layers.

### DeepRBP transcript prediction evaluation

The model effectively predicts transcript expression across both TCGA (test set) and GTEx datasets, as indicated by an averaged Spearman’s correlation of 0.854 across tumor types in TCGA and of 0.831 in GTEx, and an averaged Pearson’s correlation of and 0.979 and 0.951, respectively. Additionally, the obtained average MSE values are 0.086 for TCGA and 0.183 for GTEx, further affirming the model’s performance (see Supplementary Table 5-6 for results per tumor type).

To further assess the model performance and validate the training with the whole TCGA, we trained separate models with samples from each tumor type, showing no loss in performance. In fact, the model exhibits better generalization capabilities (see Supplementary Fig. 2-3).

Moreover, the model exhibited robustness in predicting isoform abundance, as evidenced by effectively predicting 98.5% of genes’ isoform usage in TCGA across all tumor types, and achieving a 96.7% success rate in GTEx. Detailed percentages for each tumor type can be found in Supplementary Table 7-8. For example, in liver hepatocellular carcinoma, the sum of percentages was 97.9%, with a standard deviation of 0.070.

It should be noted that the provided results on isoform usage calculation were performed only with genes whose mean expression exceeded 5 TPM across samples. The reason is the low confidence in transcript abundance computation for low-expressed genes, as mapping algorithms have access to fewer reads, resulting in more variability in the results.

#### Explainability validation with POSTAR3

We expect higher RBP-Gene scores to be associated with regulations already identified in POSTAR3. Indeed, with the considered TCGA’s acute myeloid leukemia (AML), kidney chromophobe and liver hepatocellular carcinoma RNA-seq data (see Methods), we demonstrate that this is the case. Specifically, significant differences are observed in the Mann-Whitney-Wilcoxon test (*p*.val *<* 0.05) when comparing the scores of class 0 and class 1 of POSTAR3 for each RBP in the three experiments. Supplementary Fig. 7-9 display the distribution of the obtained *p*.val for all RBPs. The scores for some selected RBPs in AML are visualized in Fig. 2A-left, distinguishing between class 0, class 1, and class NaN. For additional examples, see Supplementary Fig. 10-12 (left).

**Fig. 2.**
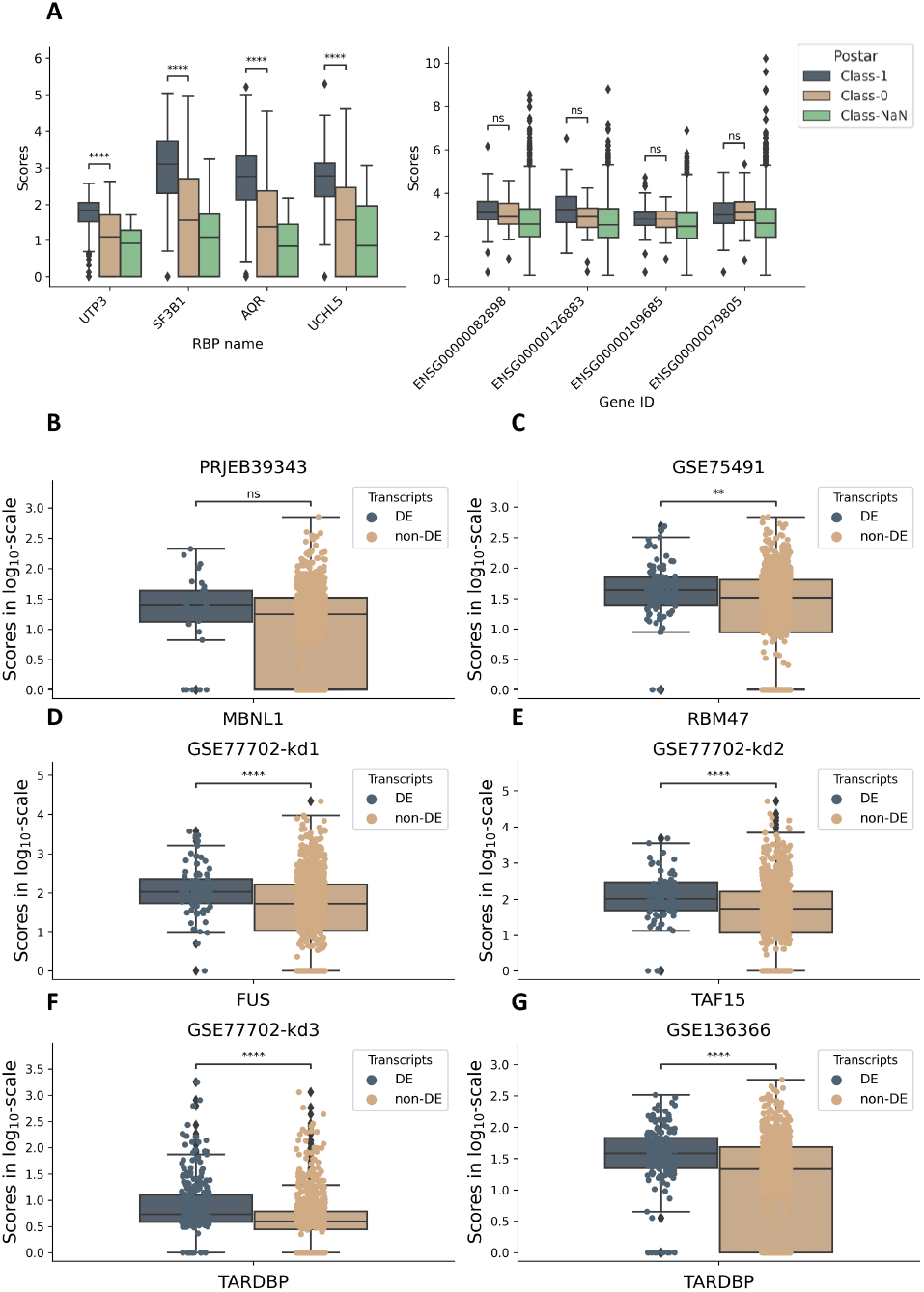
Explainability validation of DeepRBP obtained scores. **A**. Boxplot of RBP-Gene scores distinguished by class 0, 1, and N/A of POSTAR3 using acute myeloid leukemia samples across RBPs (on the left) and across genes (on the right). The 4 RBPs and genes with the highest proportion of 1s in POSTAR3 are depicted. **B-G**. RBP-Gene scores for the knockdown RBP of each of the six considered knockdown experiments.

On the other hand, performing the same analysis across the genes’ scores did not reveal significant (*p*.val *<* 0.05) differences (see Figure 2-right for selected examples in AML and Supplementary Fig. 10-12 (right) for additional ones). We believe this has to do with how DeepLIFT computes the scores, as it modifies the input of a given RBP to compute the scores of that RBP across genes, making the RBP-scores more informative. Based on this observation, in the following we focus the analyses on the RBP-scores.

To further assess the obtained scores with the proposed DeepRBP framework, we computed the AUC-ROC values for each RBP and experiment when employed to distinguish between class 0 (no regulation) and class 1 (regulation) of POSTAR3. This yielded on average AUC values of 0.7031 and standard deviation 0.05 (see Table 3). We compared these scores with alternative methods also based on DeepLIFT, showing overall the best performance in the considered POSTAR3 experiments (refer to Table 3). Additionally approaches not based on DeepLIFT to calculate *G × RBP* scores were attempted by manually setting an RBP value to 0 (knockdown) or 1 (knockup) in the predictive model, and then computing the fold-change (FC) of the predicted transcript expressions (in TPM) between two conditions: knockdown-control; knockdown-knockup; and control-knockup. Control refers to leaving the RBP unchanged. The FC of low-expressed genes (*<* 1 TPM) was set to 1 and subsequently the log_2_(FC) was calculated. The transformation to genes is done by taking the maximum value of the transcripts. These approaches also led to less-informative scores (see Supplementary Table 9).

**Table 3.** Comparison of DeepRBP explainability approach with other alternatives also based on DeepLIFT. The main difference comes from the used reference in DeepLIFT: all-zeros (kd) or the median; and the way the scores across samples are collapsed: t-statistic or sum. For each tumor type, the mean and standard deviation of the AUC calculated with the RBP scores to distinguish between classes 0 and 1 of POSTAR3 are presented.

| Tumor type | DeepRBP (Ref kd; t-stat) |  | Ref kd; sum |  | Ref median; t-stat |  | Ref median; sum |  |
| --- | --- | --- | --- | --- | --- | --- | --- | --- |
|  | mean | std | mean | std | mean | std | mean | std |
| Acute Myeloid Leukemia | 0.7147 | 0.0444 | 0.7152 | 0.0490 | 0.6513 | 0.0570 | 0.7090 | 0.0463 |
| Kidney Chromophobe | 0.7061 | 0.0594 | 0.6859 | 0.0477 | 0.6469 | 0.0512 | 0.6711 | 0.0520 |
| Liver Hep carcinoma | 0.6887 | 0.0463 | 0.7097 | 0.0519 | 0.6762 | 0.0429 | 0.7110 | 0.0426 |

Finally, we analyzed if gene scores were correlated to the number of transcripts, since RBP-Gene scores are computed by taking the maximum absolute RBP-Transcript scores of the corresponding gene’s transcripts. We obtained on average a Pearson correlation of 0.30 (Supplementary Fig. 13). Despite this statistical observation, from a biological perspective it is reasonable to expect that genes with a higher transcript count undergo more splicing events and, consequently, are regulated by a greater number of RBPs (see Supplementary Fig. 14-16).

#### Validation with RBP knockdown data

We conducted six experiments with real RBPs knockdown data (Table 2). The predictive performance of the trained DeepRBP model on both control and knockdown samples exhibited an average MSE of 0.425 and 0.488, respectively, slightly higher than with the TCGA and GTEx samples. In the same line, the average Spearman’s correlation coefficients displayed a slight reduction, with values of 0.7690 (0.7498) control (knockdown) samples (see Supplementary Table 10 for detailed results).

For ease of exposition, we focus the discussion on the TARDBP knowckdown (GSE136366 experiment). The DE analysis identified 2,564 transcripts as differentially expressed (*p*.val *<* 0.05), from the 143,561 examined. This corresponds to 191 transcripts from the list of 11,462 cancer-associated transcripts considered in this study, and to 157 distinct genes. Subsequently, DeepRBP was used to derive the explainability scores. Table 4 summarizes the results of this process for all experiments.

**Table 4.** Summary of the number of transcripts and genes differentially expressed in the real RBP knockdown experiments.

| Exp. | RBP | Num Transcripts | Num Genes |
| --- | --- | --- | --- |
| PRJEB39343 | MBNL1 | 36 | 32 |
| SRR296 | RBM47 | 114 | 95 |
| GSE77702-kd1 | FUS | 89 | 81 |
| GSE77702-kd2 | TAF15 | 87 | 80 |
| GSE77702-kd3 | TARDBP | 372 | 283 |
| GSE136366 | TARDBP | 191 | 157 |

A comparison of the RBP-Transcript scores between DE transcripts and non-DE transcripts for RBP TARDBP (GSE136366 experiment) revealed a statistically significant difference (*p*.val *<* 0.05). This trend persisted for other knockdown RBPs such as RBM47, TAF15 and TARDBP (GSE77702). Even for MBNL1, where the difference is not significant (*p*.val *>* 0.05), the median score of DE transcripts is higher than non-DE, 5.64 vs 4.43 (see Supplementary Fig. 17). Conversely, at the gene level, i.e., with the RBP-Gene scores, this disparity was found to be always statistically significant (*p*.val *<* 1 *×* 10^−2^) except for RBP MBNL1 (Fig. 2B-G).

These results showcase the capability of the proposed framework DeepRBP to identify potential RBP-Gene regulations using only control data.

#### Identifying potential RBP-Gene candidates

Thresholds to discern whether a gene is regulated by a given RBP were computed (see Methods). For instance, for RBP EFTUD2 in liver hepatocellular carcinoma the threshold is set to 1.01 (see Supplementary Fig. 18). For those RBPs not characterized in POSTAR3, since a threshold can not be computed based on the ROC curve, we take the maximum threshold among the RBPs analyzed in POSTAR3. We then select the top 50 candidates with the highest scores. In the context of liver hepatocellular carcinoma, this resulted in 303 potential candidates for RBP-Gene regulations not previously described in POSTAR3, wherein the RBP score assigned to the gene has surpassed the threshold designated for that RBP.

For example, for RBPs SRSF1 and SRSF7, known to play a key role in hepatocellular carcinoma (Shen *et al*., 2024; Lee *et al*., 2020), we identified gene ENSG00000176105 as potentially being regulated by both RBPs, and gene ENSG00000183579 by SRSF1. We also detected the following candidates of RBP-Genes which exhibited very high scores: RBP HLA-A with gene ENSG00000206503, RBP MRPL41 with gene ENSG00000163453, and RBP RBMY1A1 with gene ENSG00000164919.

Similar analysis have been performed using acute myeloid leukemia and kidney chromophobe RNA-seq data.

Potential candidates for these experiments are provided in Supplementary table 3.xlsx. Nevertheless, we acknowledge the need to further validate these results in-vitro.

## Conclusion

In this work, we presented DeepRBP, a framework to identify in-silico potential unknown splicing regulations of the form RBP-Gene. DeepRBP is based on a DL model to predict isoform usage coupled with an explainability module that computes informative RBP-Gene scores. Our experimental results demonstrate that the prediction model is capable of accurately predicting transcript expression given RBP and gene expression data. Regarding the interpretability of the network, we have shown that DeepRBP assigns significantly higher values to known RBP-Gene interactions than to RBP-Gene pairs known to have no interactions. These results have been validated using the POSTAR3 database, as well as real knockout data. Moreover, using specific thresholds for the considered RBPs, we were able to identify a handful of RBP-Gene candidates in hepatocellular carcinoma for additional in-vitro validation.

Future work includes further validation of the proposed framework with additional knockout data, and in-vitro validation of some of the identified RBP-Gene candidates, as well as increasing the number of selected genes in DeepRBP.

## Acknowledgments

We would like to thank Angel Rubio for insightful discussions. This work was partially supported by the following grants: Elkartek and RIS3 from the Basque Government [KK-2023/00001, 2023333040], Fundacion Ramon Areces, MCIN/AEI RYC2019-028578-I, Gipuzkoa Fellows (2022-FELL-000003-01), and MCIN/AEI (PID2021-126718OA-I00).

## References

Akiba, T. et al. (2019). Optuna: A next-generation hyperparameter optimization framework. In Proceedings of the 25th ACM SIGKDD international conference on knowledge discovery & data mining, pages 2623–2631.

Bergstra, J. et al. (2011). Algorithms for hyper-parameter optimization. Advances in neural information processing systems, 24.

Bergstra, J. et al. (2013). Making a science of model search: Hyperparameter optimization in hundreds of dimensions for vision architectures. In International conference on machine learning, pages 115–123. PMLR.

Carazo, F. et al. (2019). Integration of clip experiments of rna-binding proteins: a novel approach to predict context-dependent splicing factors from transcriptomic data. BMC genomics, 20, 1–11.

Chen, F. and Keleş, S. (2020). Surf: integrative analysis of a compendium of rna-seq and clip-seq datasets highlights complex governing of alternative transcriptional regulation by rna-binding proteins. Genome biology, 21, 1–23.

Cheng, S. et al. (2020). A functional network of gastric-cancer-associated splicing events controlled by dysregulated splicing factors. NAR Genomics and Bioinformatics, 2(2), lqaa013.

Dominguez, D. et al. (2018). Sequence, structure, and context preferences of human rna binding proteins. Molecular cell, 70(5), 854–867.

Ferrer-Bonsoms, J. A. et al. (2022). Eventpointer 3.0: flexible and accurate splicing analysis that includes studying the differential usage of protein-domains. NAR Genomics and Bioinformatics, 4(3), lqac067.

Gárate-Rascón, M. et al. (2022). Slu7: A new hub of gene expression regulation—from epigenetics to protein stability in health and disease. International Journal of Molecular Sciences, 23(21), 13411.

Goldman, M. J. et al. (2020). Visualizing and interpreting cancer genomics data via the xena platform. Nature biotechnology, 38(6), 675–678.

Hong, S. (2017). Rna binding protein as an emerging therapeutic target for cancer prevention and treatment. Journal of cancer prevention, 22(4), 203.

Ioffe, S. and Szegedy, C. (2015). Batch normalization: Accelerating deep network training by reducing internal covariate shift. arXiv preprint arXiv:1502.03167 .

Kapeli, K. et al. (2016). Distinct and shared functions of als-associated proteins tdp-43, fus and taf15 revealed by multisystem analyses. Nature communications, 7(1), 12143.

Law, C. W. et al. (2014). voom: Precision weights unlock linear model analysis tools for rna-seq read counts. Genome biology, 15, 1–17.

Lee, S. E. et al. (2020). Alternative splicing in hepatocellular carcinoma. Cellular and Molecular Gastroenterology and Hepatology, 10(4), 699–712.

Li, J. et al. (2021). Alternative splicing perturbation landscape identifies rna binding proteins as potential therapeutic targets in cancer. Molecular Therapy-Nucleic Acids, 24, 792–806.

Lonsdale, J. et al. (2013). The genotype-tissue expression (gtex) project. Nature genetics, 45(6), 580–585.

Oltean, S. and Bates, D. O. (2014). Hallmarks of alternative splicing in cancer. Oncogene, 33(46), 5311–5318.

Park, J. W. et al. (2016). rmaps: Rna map analysis and plotting server for alternative exon regulation. Nucleic acids research, 44(W1), W333–W338.

Ramanathan, M. et al. (2019). Methods to study rna-protein interactions. Nature methods, 16(3), 225–234.

Roczniak-Ferguson, A. and Ferguson, S. M. (2019). Pleiotropic requirements for human tdp-43 in the regulation of cell and organelle homeostasis. Life Science Alliance, 2(5).

Sebestyén, E. et al. (2016). Large-scale analysis of genome and transcriptome alterations in multiple tumors unveils novel cancer-relevant splicing networks. Genome research, 26(6), 732–744.

Shen, S. et al. (2014). rmats: robust and flexible detection of differential alternative splicing from replicate rna-seq data. Proceedings of the National Academy of Sciences, 111(51), E5593–E5601.

Shen, W. et al. (2024). Srsf7 is a promising prognostic biomarker in hepatocellular carcinoma and is associated with immune infiltration. Genes & Genomics, 46(1), 49–64.

Shrikumar, A. et al. (2017). Learning important features through propagating activation differences. In International conference on machine learning, pages 3145–3153. PMLR.

Sondka, Z. et al. (2018). The cosmic cancer gene census: describing genetic dysfunction across all human cancers. Nature Reviews Cancer, 18(11), 696–705.

Warren, A. et al. (2021). Global computational alignment of tumor and cell line transcriptional profiles. Nature communications, 12(1), 22.

Weinstein, J. N. et al. (2013). The cancer genome atlas pan-cancer analysis project. Nature genetics, 45(10), 1113–1120.

Yang, Y. et al. (2016). Determination of a comprehensive alternative splicing regulatory network and combinatorial regulation by key factors during the epithelial-to-mesenchymal transition. Molecular and cellular biology, 36(11), 1704–1719.

Zhang, Z. et al. (2019). Deep-learning augmented rna-seq analysis of transcript splicing. Nature methods, 16(4), 307–310.

Zhao, W. et al. (2022). Postar3: an updated platform for exploring post-transcriptional regulation coordinated by rna-binding proteins. Nucleic Acids Research, 50(D1), D287–D294.

