## Supplementary Material for "DeepRBP: A novel deep neural network for inferring splicing regulation"

### Supplementary data for manuscript "DeepRBP: A novel deep neural network for inferring splicing regulation"

#### Supplementary Figures

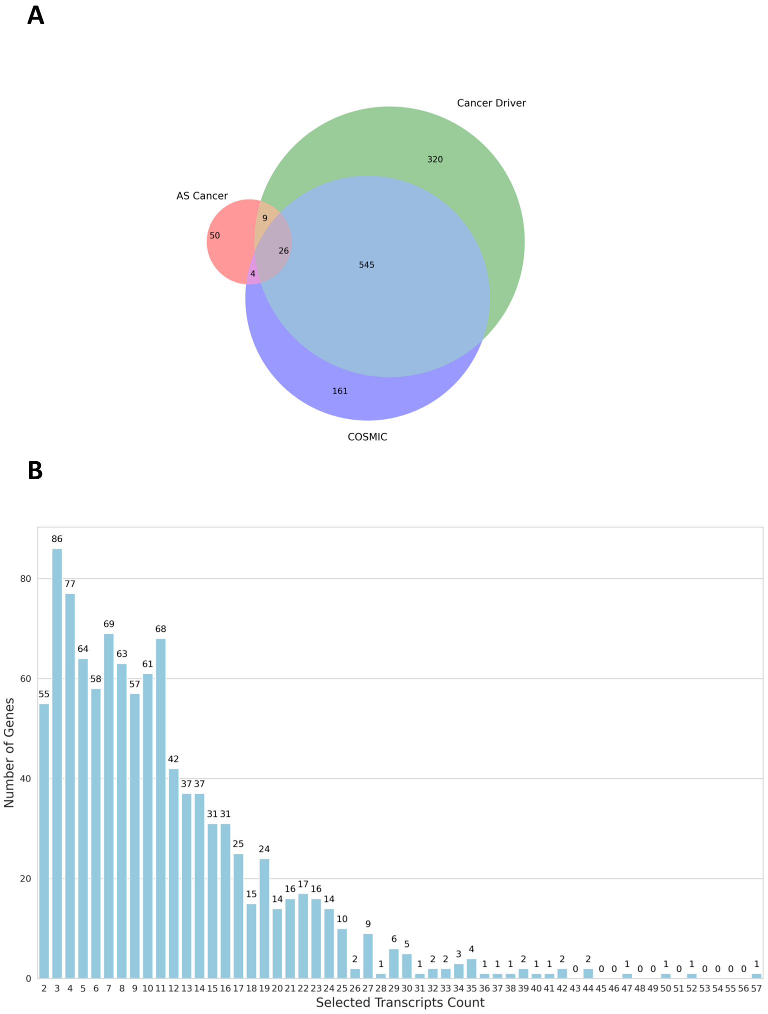


Figure 1: **A.** Venn Diagram of the genes considered in this study. We compiled a list of protein-coding genes related to splicing and cancer by integrating data from various sources, including AS Cancer genes, Cancer driver genes, and COSMIC genes. Only genes with more than one transcript were included. **B.** Histogram of the number of transcripts per gene.

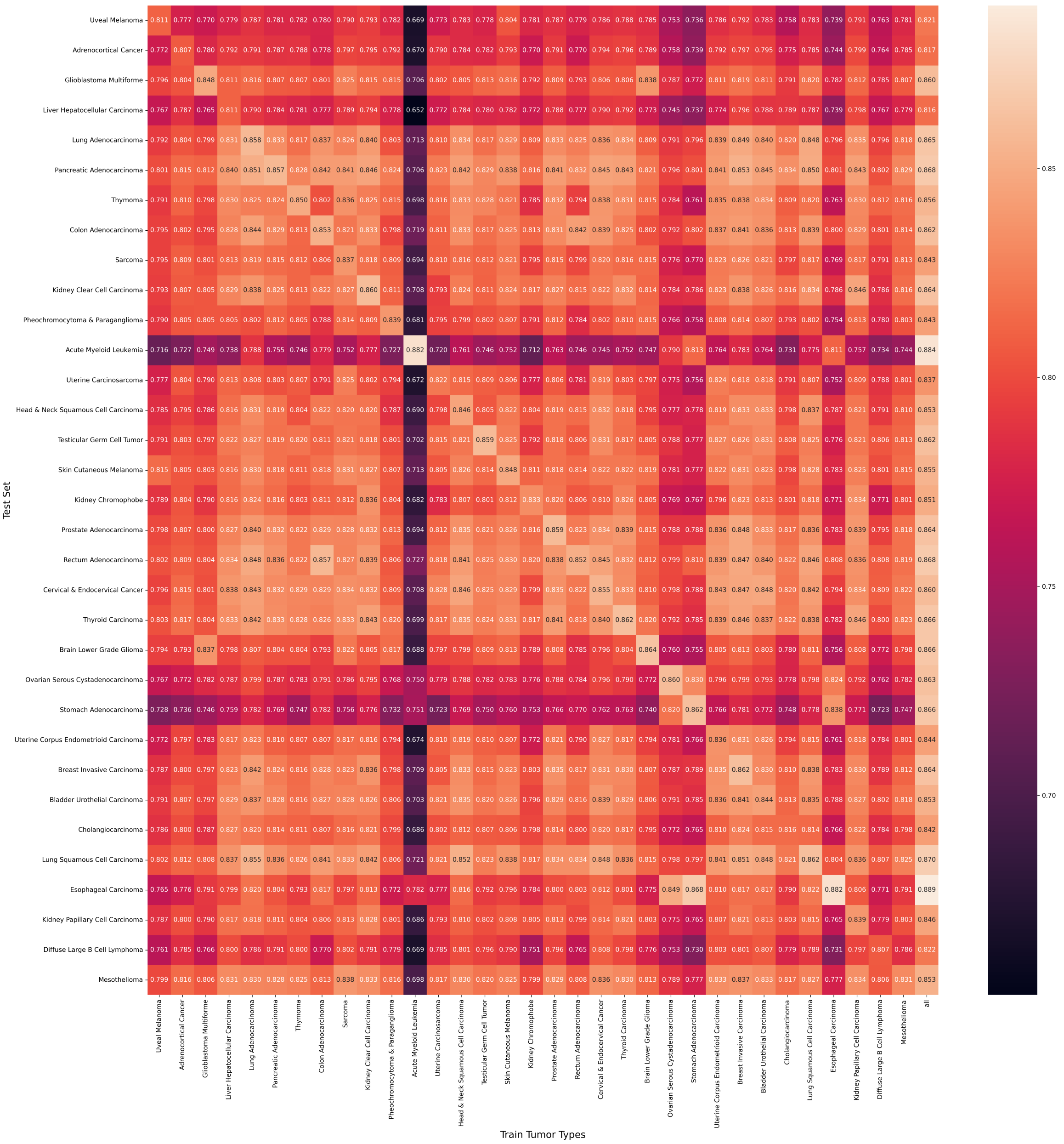


Figure 2: Spearman correlation of models trained with different tumor types individually versus using all tumor types in the test sets. The columns display the samples used to train the model, and the rows show the name of the test set.

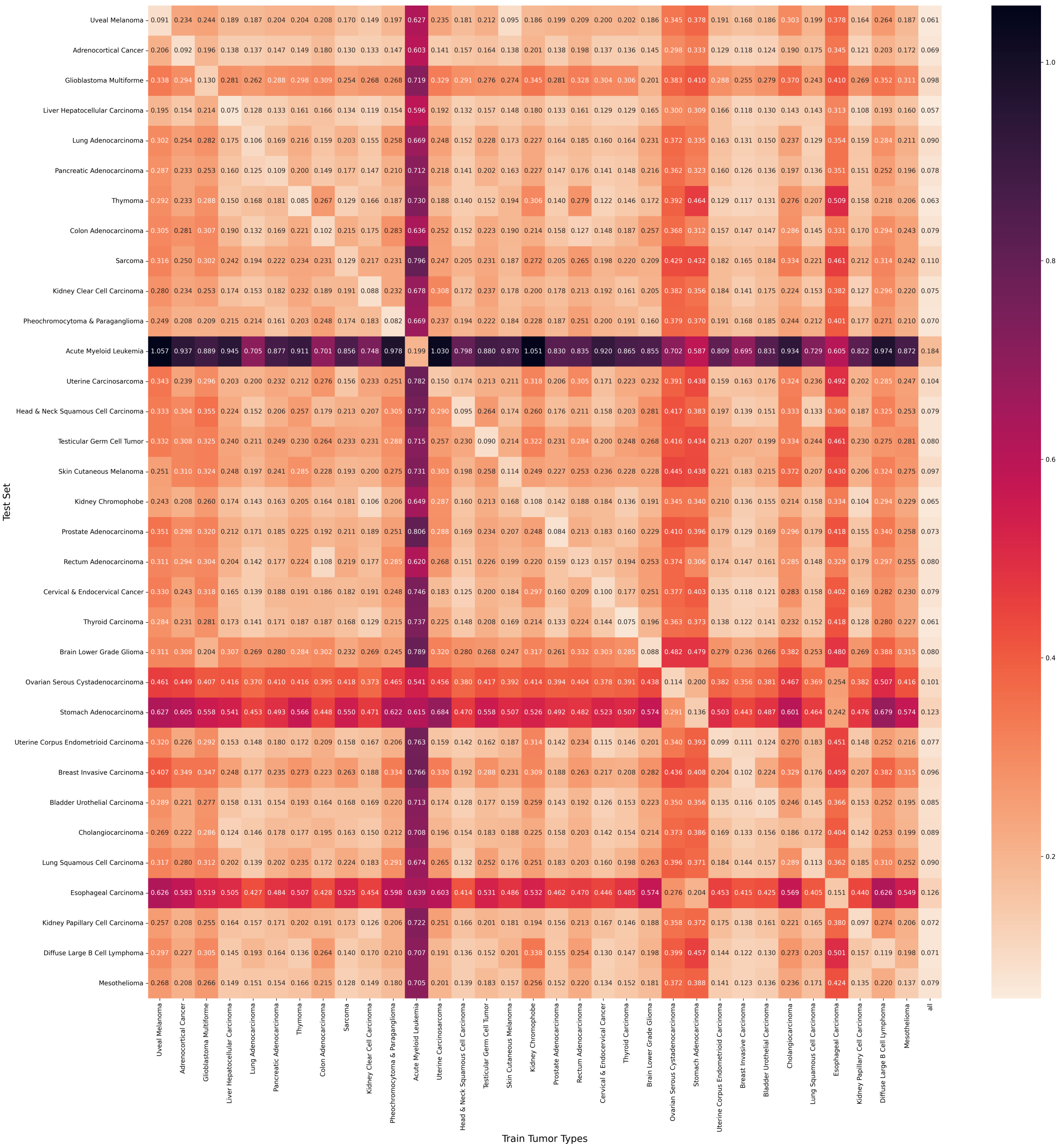


Figure 3: MSE of models trained with different tumor types individually versus using all tumor types in the test sets. The columns display the samples used to train the model, and the rows show the name of the test set.

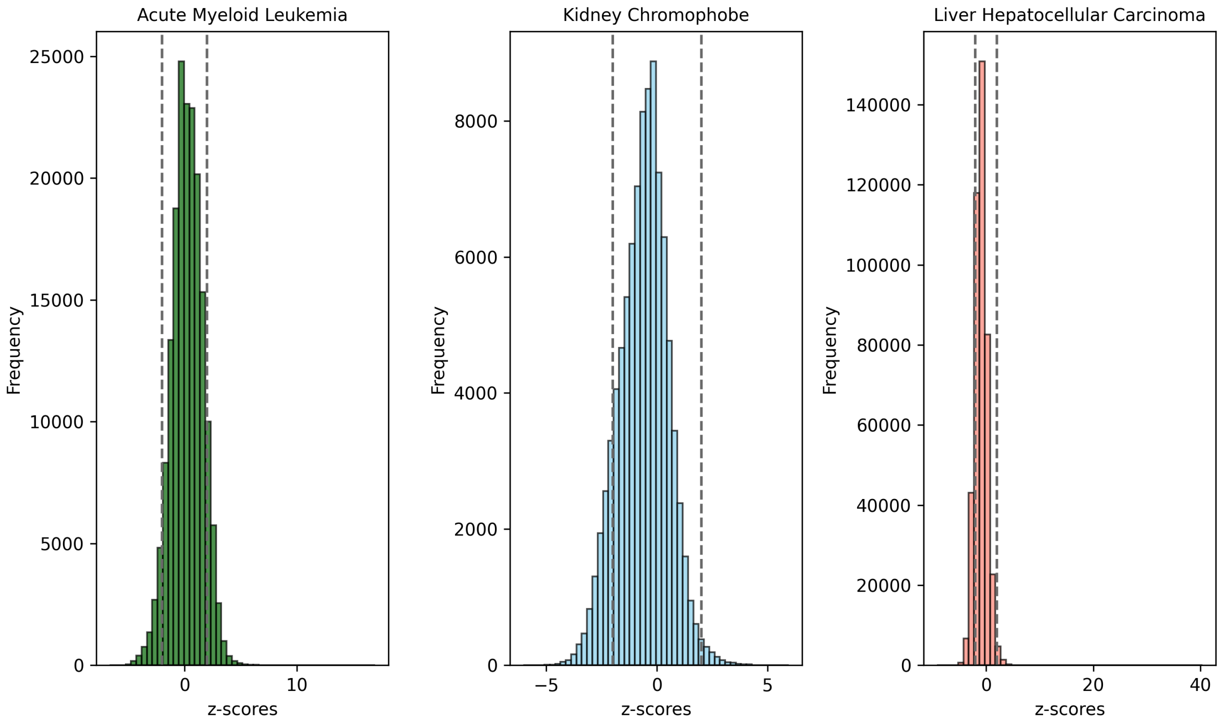


Figure 4: Distribution of RBP z-scores in acute myeloid leukemia, kidney chromophobe and liver hepatocellular carcinoma TCGA tumor samples before truncating scores to $(-2, +2)$ and displacing values to the range zero to one.

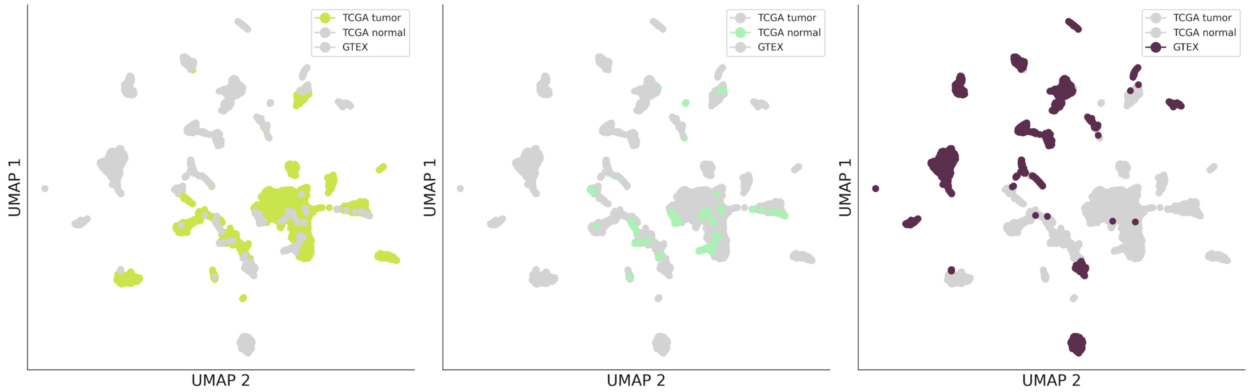


Figure 5: UMAP embedding of the expression (in log2(TPM+1)) of TCGA and GTEx samples across three conditions: TCGA tumor samples (highlighted in pistachio), TCGA normal samples (highlighted in light green), and GTEx samples (highlighted in purple), with the remaining samples depicted in shaded gray for comparison.

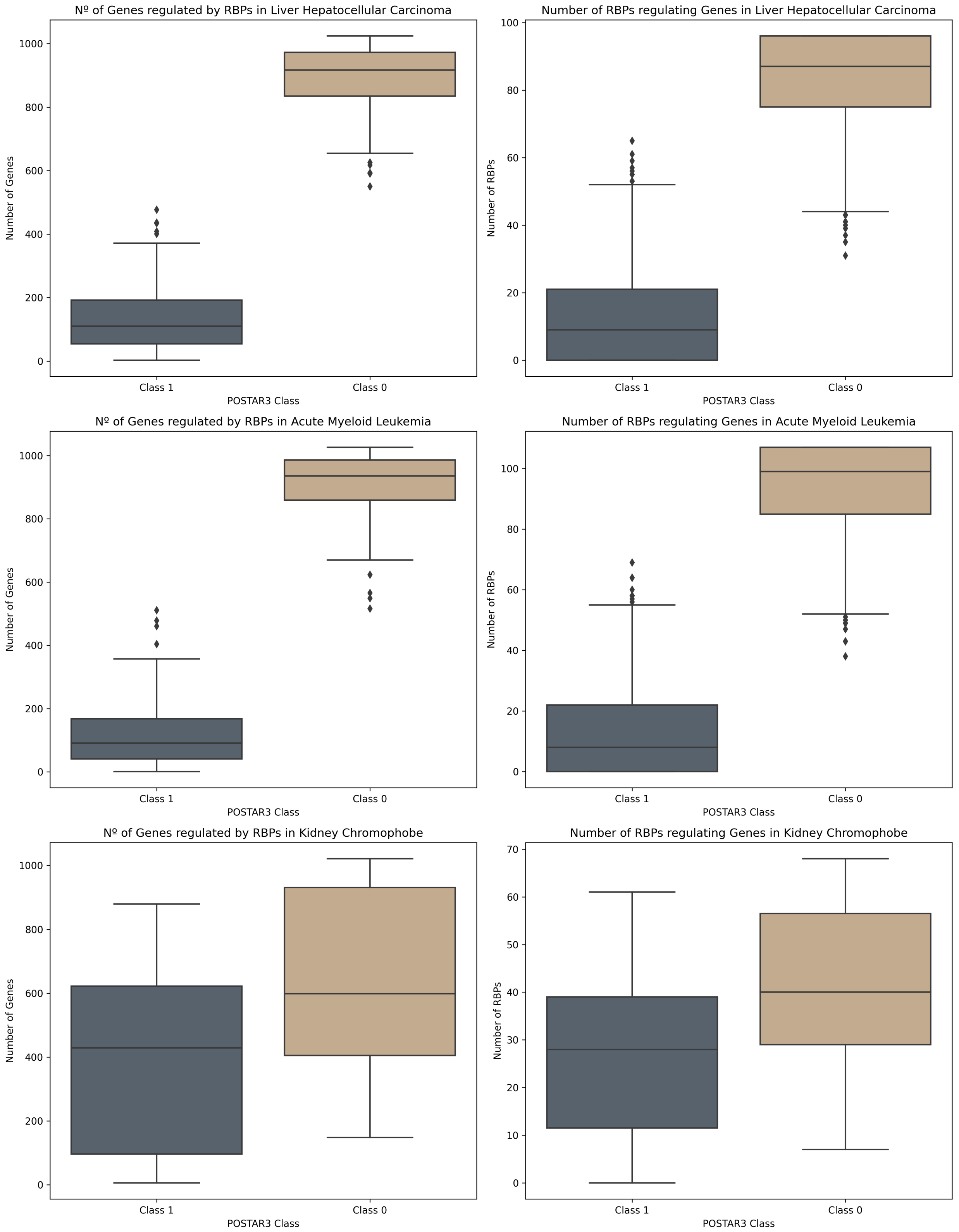


Figure 6: Boxplots of the number of genes regulated by a given RBP (left) and number of RBPs regulating a given gene (right) for class 1 and class 0 samples for the three considered POSTAR3 experiments.

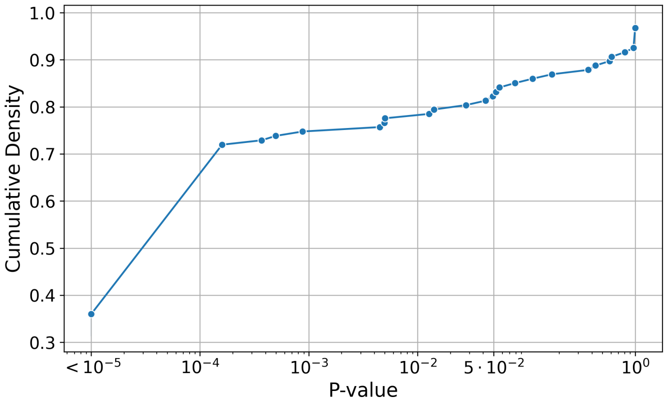


Figure 7: Cumulative distribution of p-values obtained from the Mann-Whitney-Wilcoxon test corrected by Bonferroni method for distinguishing scores between class 0 and class 1 using samples from acute myeloid leukemia.

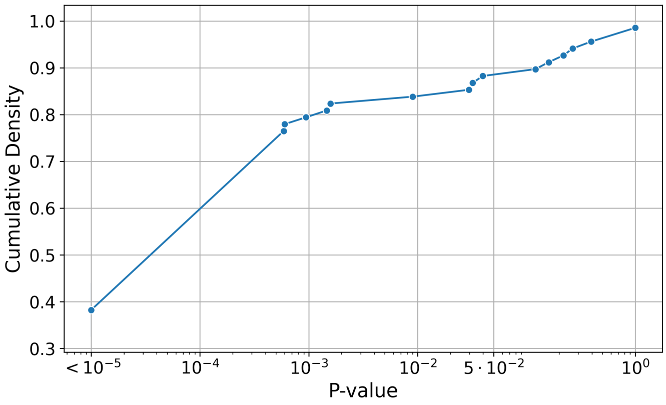


Figure 8: Cumulative distribution of p-values obtained from the Mann-Whitney-Wilcoxon test corrected by Bonferroni method for distinguishing scores between class 0 and class 1 using samples from kidney chromophobe.

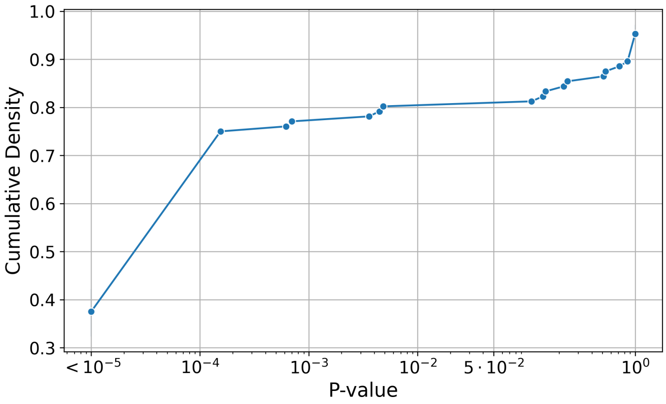


Figure 9: Cumulative distribution of p-values obtained from the Mann-Whitney-Wilcoxon test corrected by Bonferroni method for distinguishing scores between class 0 and class 1 using samples from kidney chromophobe.

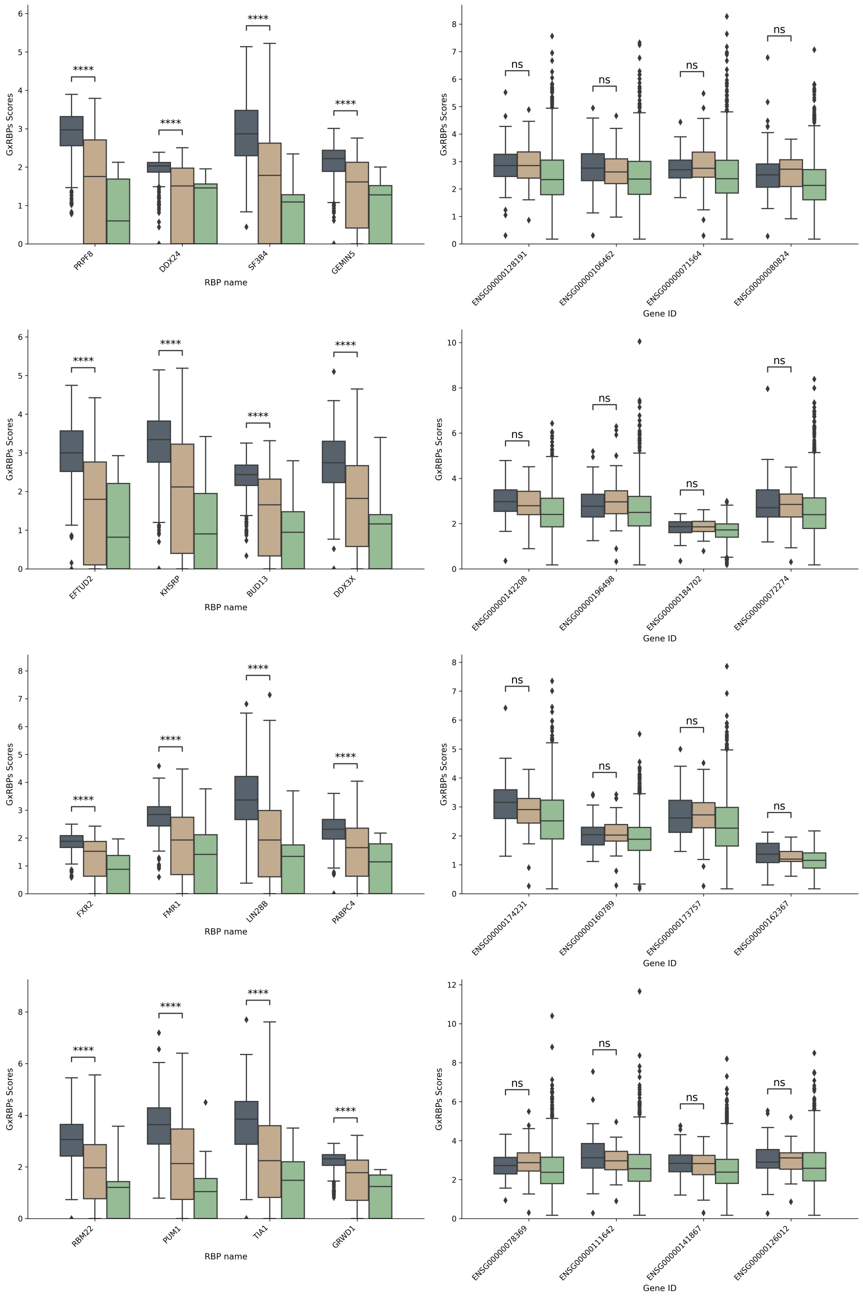


Figure 10: Boxplot of RBP-Gene scores by class (0, 1, and N/A) in POSTAR3 for the Acute Myeloid Leukemia samples, depicted by RBP (**left**) and by gene (**right**). The selected RBPs and Genes are those exhibiting the largest number of 1s in POSTAR3.

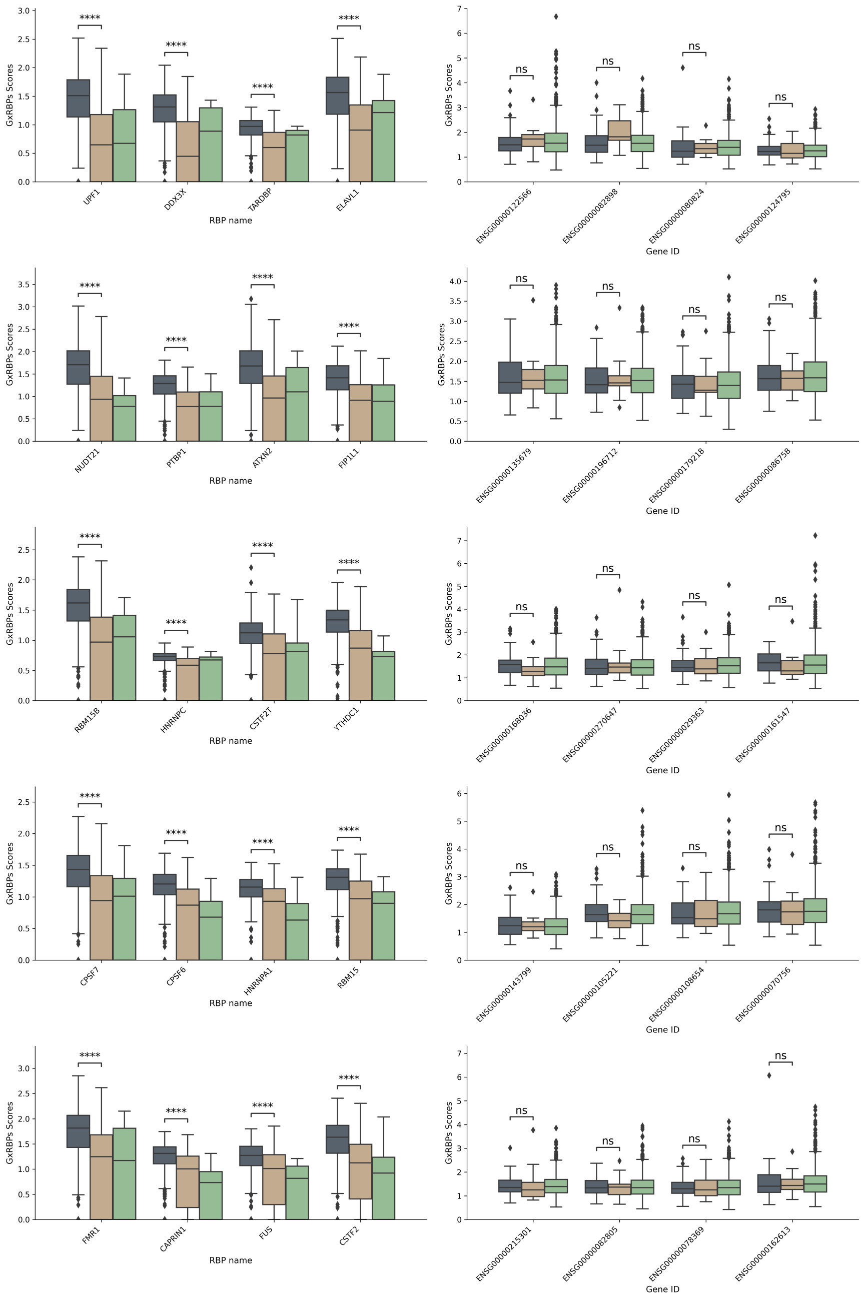


Figure 11: Boxplot of RBP-Gene scores by class (0, 1, and N/A) in POSTAR3 for the Kidney Chromophobe samples, depicted by RBP (**left**) and by gene (**right**). The selected RBPs and Genes are those exhibiting the largest number of 1s in POSTAR3.

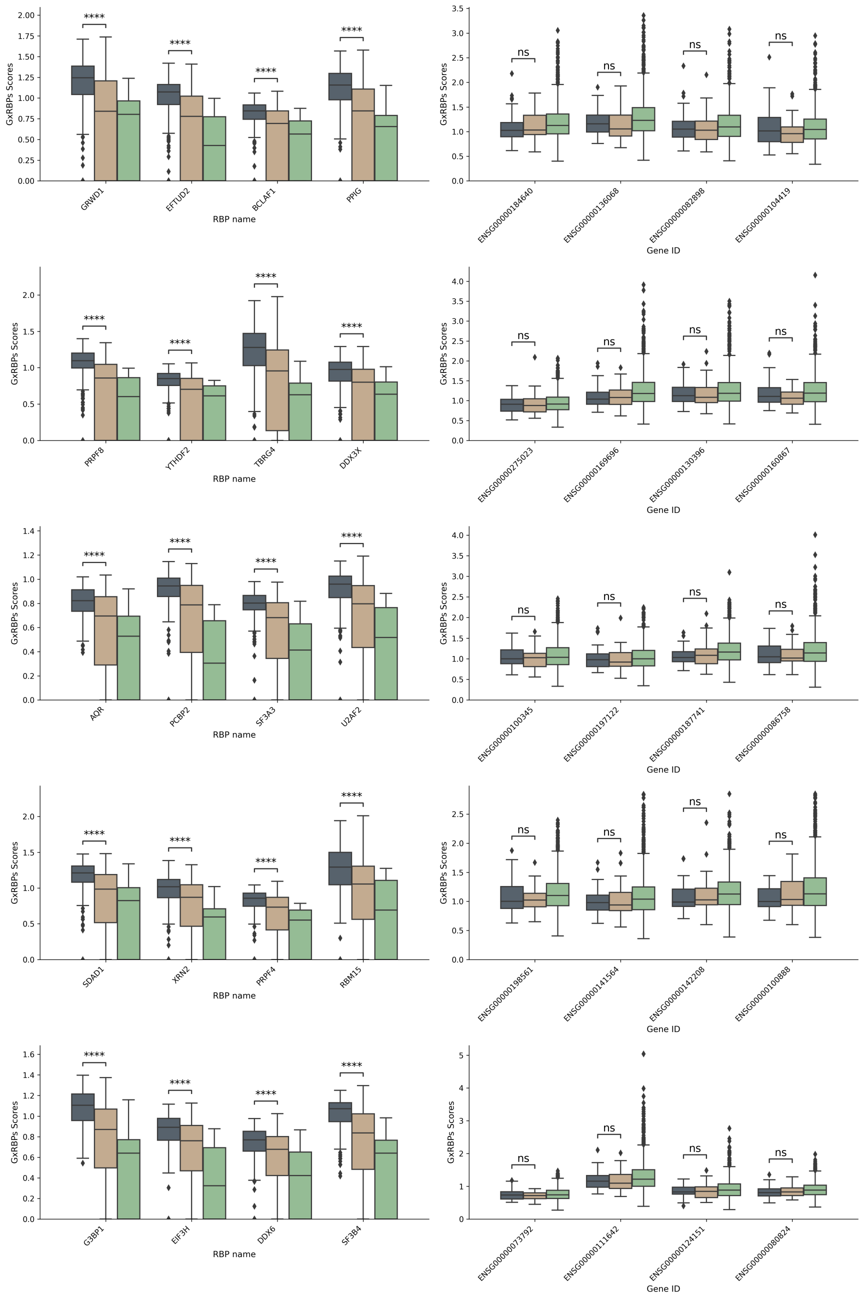


Figure 12: Boxplot of RBP-Gene scores by class (0, 1, and N/A) in POSTAR3 for the Liver Hepatocellular Carcinoma samples, depicted by RBP (**left**) and by gene (**right**). The selected RBPs and Genes are those exhibiting the largest number of 1s in POSTAR3.

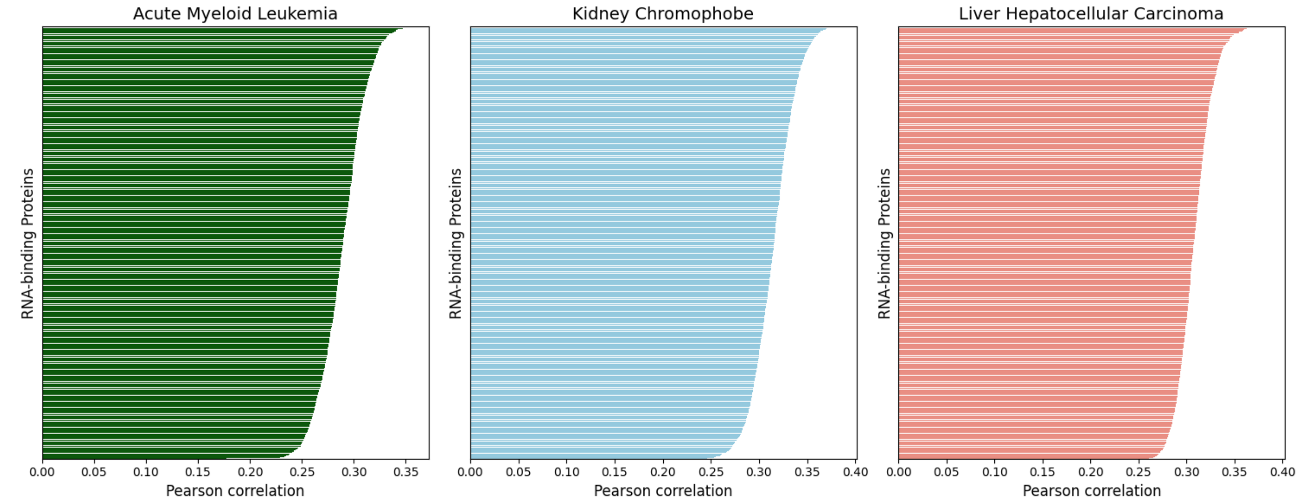


Figure 13: Pearson correlation between the gene scores computed by DeepRBP and the number of transcripts per gene for the 1,282 RBPs (sorted by correlation value) for the three considered POSTAR3 experiments. The results show a mean correlation of 0.2892 for acute myeloid leukemia, 0.3152 for kidney chromophobe, and 0.3086 for liver hepatocellular carcinoma.

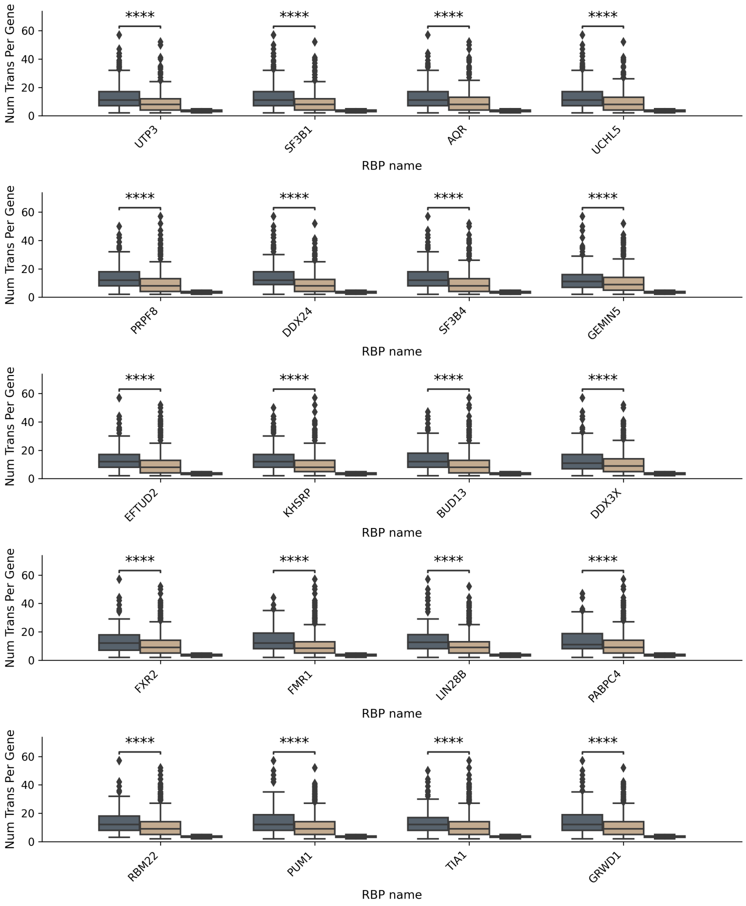


Figure 14: Number of transcripts for class 1 and class 0 RBP-Gene pairs in POSTAR3 for the K562 cell-line, for RBPs with most validated values.

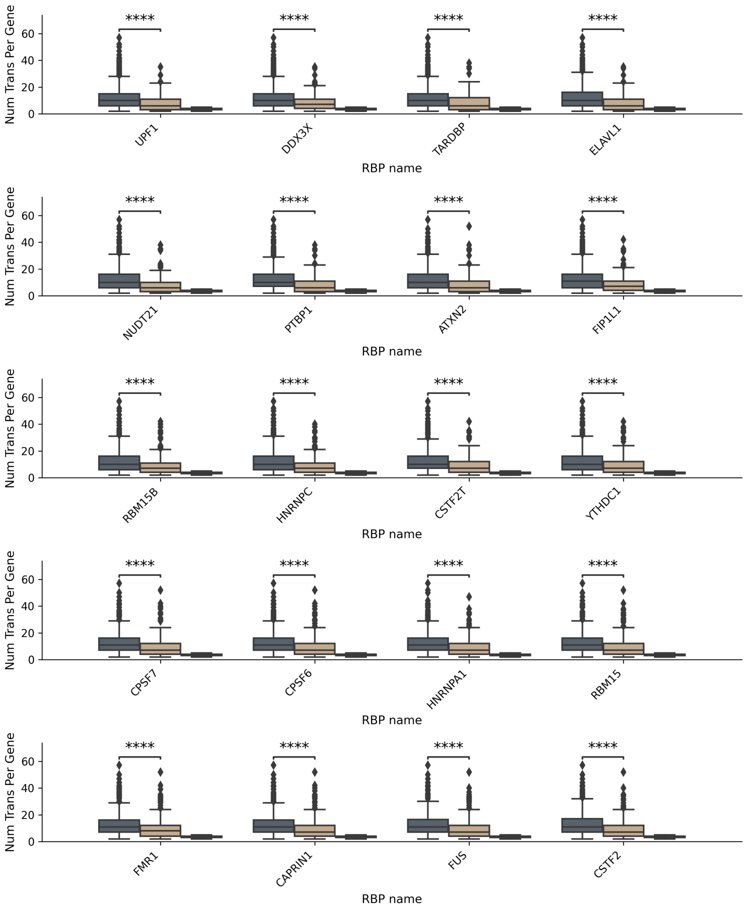


Figure 15: Number of transcripts for class 1 and class 0 RBP-Gene pairs in POSTAR3 for the HEK293 cell-line, for RBPs with most validated values.

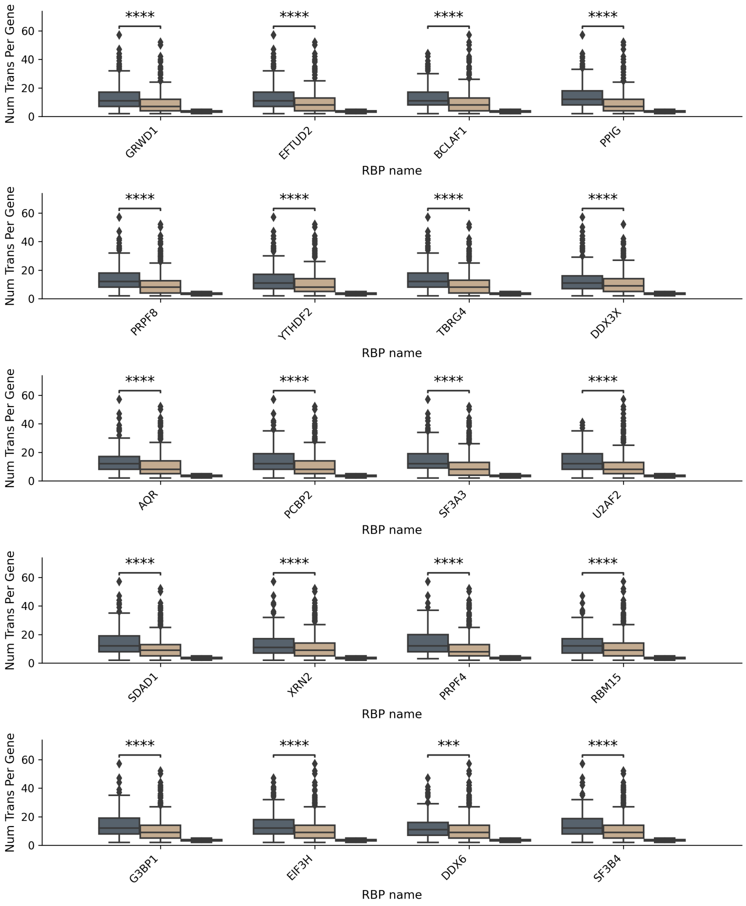


Figure 16: Number of transcripts for class 1 and class 0 RBP-Gene pairs in POSTAR3 for the huh7 and HepG2 cell-line, for RBPs with most validated values.

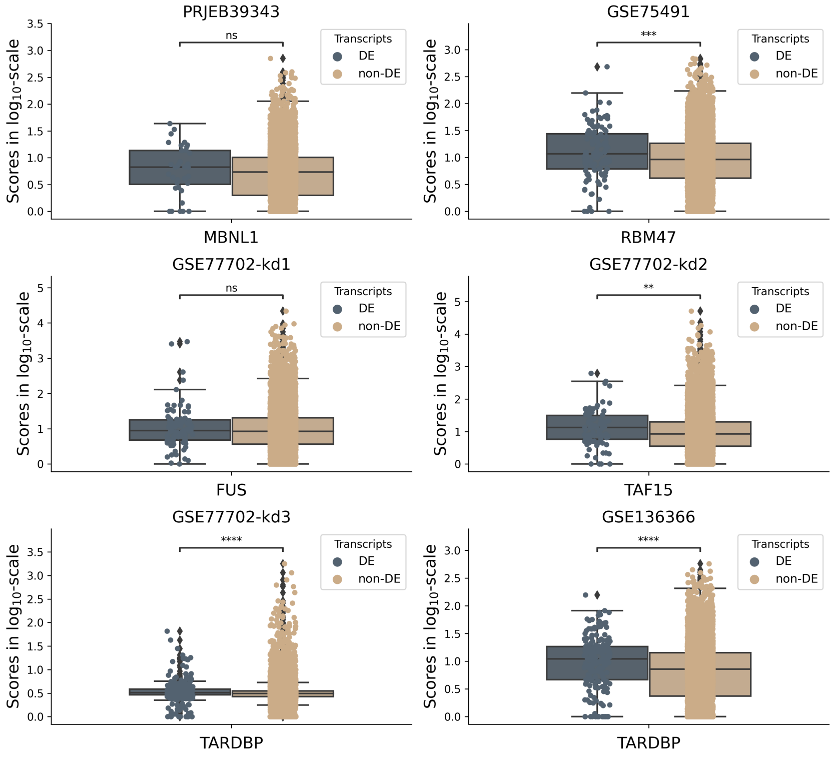


Figure 17: Boxplot of RBP-Transcript scores for the knockdowned RBP of each of the six considered experiments, for differentially expressed (DE) and non-DE transcripts.

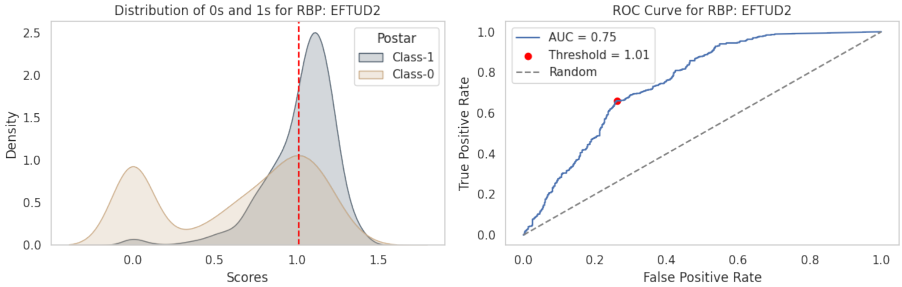


Figure 18: **Left:** Distribution of class 0 and class 1 RBP-Gene scores for RBP EFTUD2 obtained with the liver hepatocellular carcinoma samples (left). **Right:** ROC curve for the same scores, with the computed threshold for RBP EFTUD2.

#### Supplementary Tables

Table 1: Number of samples used for training and validation in the model optimization with Optuna for each tumor type.

| Tissue | N train | N validation |
|---|---|---|
| Acute myeloid leukemia | 44 | 11 |
| Uterine carcinosarcoma | 14 | 4 |
| Pheochromocytoma & paraganglioma | 47 | 12 |
| Testicular germ cell tumor | 39 | 10 |
| Thyroid carcinoma | 145 | 37 |
| Lung adenocarcinoma | 147 | 37 |
| Mesothelioma | 22 | 6 |
| Adrenocortical cancer | 19 | 5 |
| Gliobastoma multiforme | 43 | 11 |
| Diffuse large B cell lymphoma | 12 | 3 |
| Thymoma | 30 | 8 |
| Stomach adenocarcinoma | 115 | 29 |
| Head & neck squamous cell carcinoma | 144 | 36 |
| Liver hepatocelullar carcinoma | 107 | 27 |
| Pancreatic adenocarcinoma | 46 | 12 |
| Ovarian serous cystadenocarcinoma | 108 | 28 |
| Lung squamous cell carcinoma | 140 | 35 |
| Sarcoma | 67 | 17 |
| Kidney chromophobe | 23 | 6 |
| Colon adenocarcinoma | 84 | 22 |
| Cholangiocarcinoma | 11 | 3 |
| Brain lower grade glioma | 133 | 34 |
| Esophageal carcinoma | 49 | 13 |
| Bladder urothelial carcinoma | 108 | 28 |
| Cervical & endocervical cancer | 79 | 20 |
| Uterine corpus endometrioid carcinoma | 52 | 13 |
| Prostate adenocarcinoma | 140 | 35 |
| Kidney papillary cell carcinoma | 81 | 21 |
| Kidney clear cell carcinoma | 154 | 39 |
| Uveal melanoma | 20 | 5 |
| Skin cutaneous melanoma | 120 | 30 |
| Rectum adenocarcinoma | 26 | 7 |
| Breast invasive carcinoma | 309 | 78 |

Table 2: POSTAR3 experiment details: created GxRBP dataset name along with the used cell-line, number of RBPs, and the proportion of class 1 samples. The number of genes matching those in DeepRBP is 1027 in all cases.

| Name Dataset | Num. RBPs | % of 1s | Cell lines |
|---|---|---|---|
| Brain general GxRBP | 5 | 31.88 | frontal cortex, hippocampus |
| Skin GxRBP | 1 | 9.93 | SK-MEL-103 |
| Mieloid GxRBP | 120 | 12.23 | K562 |
| Bone Marrow GxRBP | 3 | 19.21 | SH-SY5Y |
| Pancreas GxRBP | 1 | 10.52 | Panc1, PL45 |
| Prostate carcinoma GxRBP | 1 | 37.78 | LAPC4, DU145, LNCaP |
| Myocardium GxRBP | 1 | 73.52 | Cardiac |
| Ovarian GxRBP | 1 | 8.57 | A2780 |
| T cell GxRBP | 1 | 74.59 | T cell |
| Adrenal gland GxRBP | 2 | 15.14 | adrenal gland |
| Liver all GxRBP | 107 | 12.87 | Huh7, HepG2 |
| Kidney embryo GxRBP | 81 | 36.95 | HEK293T, HEK293, HEK293-FRT, HEK293T-no-SRRM4<br>TREX-FLP-in-293T cells, HEK293T-plus-SRRM4 |
| Brain gliobastoma GxRBP | 2 | 37.24 | MGG8, T98G |
| Prostate adenocarcinoma GxRBP | 1 | 28.04 | PC3 |
| Breast adenocarcinoma GxRBP | 2 | 9.40 | MDA-MB-231, MDA-LM2 |
| Uterus Cervix GxRBP | 30 | 25.54 | HeLa, HeLa-treated with puromycoin |
| Kidney papilloma GxRBP | 1 | 2.53 | HK-2 |

Table 3: Top 5 models with less MSE value in Optuna optimization of hyperparameters

| Model | Activations functions | Learning rates | Hidden layers number | N nodes first layer | Node number scaler | N nodes constant | MSE val (obj. func) |
|---|---|---|---|---|---|---|---|
| # 87 | ReLU | 0.0001 | 3 | 1024 | 8 | True | 0.0994 |
| # 61 | Tanh | 0.01 | 2 | 1024 | 8 | - | 0.1004 |
| # 48 | ReLU | 0.01 | 3 | 1024 | 4 | False | 0.1015 |
| # 58 | ReLU | 0.001 | 3 | 512 | 8 | True | 0.1017 |
| **# 41** | **ReLU** | **0.001** | **2** | **1024** | **8** | - | **0.1031** |

Table 4: Top 5 models with greatest Spearman's value in Optuna optimization of hyperparameters

| Model | Activations functions | Learning rates | Hidden layers number | N nodes first layer | Node number scaler | N nodes constant | Spearman's corr. Val |
|---|---|---|---|---|---|---|---|
| # 87 | ReLU | 0.0001 | 3 | 1024 | 8 | True | 0.8607 |
| # 94 | Sigmoid | 0.001 | 3 | 1024 | 4 | True | 0.8605 |
| # 19 | ReLU | 0.001 | 2 | 2048 | 8 | - | 0.8604 |
| # 80 | Sigmoid | 0.01 | 3 | 1024 | 4 | True | 0.8601 |
| **# 41** | **ReLU** | **0.001** | **2** | **1024** | **8** | - | **0.8600** |

Table 5: Pearson correlation, spearman's correlation and mean squared error obtained using Deep-RBP predictor on TCGA test samples per tumor type.

| Tissue | N | Pearson corr | Spearman's corr | MSE |
|---|---|---|---|---|
| Acute myeloid leukemia | 35 | 0.962 | 0.884 | 0.184 |
| Uterine carcinosarcoma | 12 | 0.975 | 0.837 | 0.104 |
| Pheochromocytoma & paraganglioma | 37 | 0.980 | 0.843 | 0.070 |
| Testicular germ cell tumor | 31 | 0.981 | 0.862 | 0.080 |
| Thyroid carcinoma | 279 | 0.985 | 0.866 | 0.061 |
| Lung adenocarcinoma | 115 | 0.978 | 0.865 | 0.090 |
| Mesothelioma | 18 | 0.981 | 0.853 | 0.079 |
| Adrenocortical cancer | 16 | 0.979 | 0.817 | 0.069 |
| Gliobastoma multiforme | 35 | 0.977 | 0.860 | 0.098 |
| Diffuse large B cell lymphoma | 10 | 0.981 | 0.822 | 0.071 |
| Thymoma | 25 | 0.985 | 0.856 | 0.063 |
| Stomach adenocarcinoma | 90 | 0.969 | 0.866 | 0.123 |
| Head & neck squamous cell carcinoma | 113 | 0.981 | 0.853 | 0.079 |
| Liver hepatocelullar carcinoma | 85 | 0.980 | 0.816 | 0.057 |
| Pancreatic adenocarcinoma | 37 | 0.981 | 0.868 | 0.078 |
| Ovarian serous cystadenocarcinoma | 86 | 0.974 | 0.863 | 0.101 |
| Lung squamous cell carcinoma | 110 | 0.979 | 0.870 | 0.090 |
| Sarcoma | 53 | 0.975 | 0.843 | 0.110 |
| Kidney chromophobe | 19 | 0.981 | 0.851 | 0.065 |
| Colon adenocarcinoma | 67 | 0.981 | 0.862 | 0.079 |
| Cholangiocarcinoma | 9 | 0.977 | 0.842 | 0.089 |
| Brain lower grade glioma | 105 | 0.981 | 0.866 | 0.080 |
| Esophageal carcinoma | 39 | 0.972 | 0.889 | 0.126 |
| Bladder urothelial carcinoma | 86 | 0.979 | 0.853 | 0.085 |
| Cervical & endocervical cancer | 62 | 0.981 | 0.860 | 0.079 |
| Uterine corpus endometrioid carcinoma | 41 | 0.980 | 0.844 | 0.077 |
| Prostate adenocarcinoma | 110 | 0.983 | 0.864 | 0.073 |
| Kidney papillary cell carcinoma | 65 | 0.981 | 0.846 | 0.072 |
| Kidney clear cell carcinoma | 121 | 0.981 | 0.864 | 0.075 |
| Uveal melanoma | 16 | 0.982 | 0.821 | 0.061 |
| Skin cutaneous melanoma | 94 | 0.977 | 0.855 | 0.097 |
| Rectum adenocarcinoma | 21 | 0.981 | 0.868 | 0.080 |
| Breast invasive carcinoma | 243 | 0.978 | 0.864 | 0.096 |

Table 6: Pearson correlation, spearman's correlation and mean squared error obtained using Deep-RBP predictor on GTeX samples per tissue.

| Tissue | N | Pearson corr | Spearman's corr | MSE |
|---|---|---|---|---|
| Vagina | 85 | 0.955 | 0.851 | 0.195 |
| Uterus | 78 | 0.957 | 0.852 | 0.202 |
| Kidney cortex | 28 | 0.955 | 0.823 | 0.127 |
| Heart left ventricle | 202 | 0.937 | 0.786 | 0.143 |
| Cells ebv transformed lymphocytes | 107 | 0.954 | 0.852 | 0.213 |
| Fallopian tube | 5 | 0.957 | 0.851 | 0.196 |
| Pancreas | 167 | 0.954 | 0.811 | 0.097 |
| Whole blood | 337 | 0.939 | 0.767 | 0.137 |
| Heart atrial appendage | 175 | 0.938 | 0.814 | 0.178 |
| Prostate | 100 | 0.960 | 0.851 | 0.170 |
| Nerve tibial | 278 | 0.953 | 0.846 | 0.219 |
| Ovary | 88 | 0.955 | 0.844 | 0.196 |
| Pituitary | 107 | 0.957 | 0.846 | 0.167 |
| Breast mammary tissue | 243 | 0.958 | 0.851 | 0.175 |
| Lung | 287 | 0.952 | 0.852 | 0.226 |
| Testis | 165 | 0.890 | 0.787 | 0.424 |
| Cells transformed fibroblasts | 256 | 0.949 | 0.833 | 0.230 |
| Spleen | 100 | 0.960 | 0.848 | 0.169 |
| Stomach | 175 | 0.957 | 0.831 | 0.134 |
| Small intestine terminal ileum | 92 | 0.960 | 0.854 | 0.158 |
| Liver | 110 | 0.956 | 0.803 | 0.109 |
| Bladder | 9 | 0.956 | 0.846 | 0.196 |
| Muscle skeletal | 396 | 0.936 | 0.778 | 0.194 |
| Minor salivary gland | 55 | 0.958 | 0.852 | 0.154 |
| Adrenal gland | 128 | 0.959 | 0.834 | 0.141 |
| Thyroid | 279 | 0.956 | 0.851 | 0.199 |

Table 7: Mean and standard deviation of overall predicted transcript abundance per gene (percentage of genes directed towards each transcript), considering genes with mean expression exceeding 5 TPMs in the TCGA dataset.

| Tissue | N genes | Mean Percentages | Std Percentages |
|---|---|---|---|
| Acute myeloid leukemia | 741 | 0.948 | 0.084 |
| Uterine carcinosarcoma | 742 | 0.992 | 0.078 |
| Pheochromocytoma & paraganglioma | 671 | 0.982 | 0.064 |
| Testicular germ cell tumor | 761 | 0.987 | 0.075 |
| Thyroid carcinoma | 740 | 0.996 | 0.059 |
| Lung adenocarcinoma | 794 | 0.986 | 0.061 |
| Mesothelioma | 743 | 0.988 | 0.069 |
| Adrenocortical cancer | 639 | 0.971 | 0.080 |
| Gliobastoma multiforme | 754 | 0.997 | 0.063 |
| Diffuse large B cell lymphoma | 640 | 0.985 | 0.087 |
| Thymoma | 764 | 0.995 | 0.063 |
| Stomach adenocarcinoma | 809 | 0.969 | 0.082 |
| Head & neck squamous cell carcinoma | 750 | 0.988 | 0.064 |
| Liver hepatocelullar carcinoma | 590 | 0.979 | 0.070 |
| Pancreatic adenocarcinoma | 787 | 0.989 | 0.057 |
| Ovarian serous cystadenocarcinoma | 733 | 0.974 | 0.078 |
| Lung squamous cell carcinoma | 793 | 0.984 | 0.063 |
| Sarcoma | 799 | 0.992 | 0.076 |
| Kidney chromophobe | 724 | 0.971 | 0.064 |
| Colon adenocarcinoma | 739 | 0.993 | 0.058 |
| Cholangiocarcinoma | 721 | 0.986 | 0.065 |
| Brain lower grade glioma | 773 | 0.984 | 0.057 |
| Esophageal carcinoma | 819 | 0.969 | 0.069 |
| Bladder urothelial carcinoma | 751 | 0.986 | 0.068 |
| Cervical & endocervical cancer | 751 | 0.998 | 0.067 |
| Uterine corpus endometrioid carcinoma | 734 | 0.988 | 0.073 |
| Prostate adenocarcinoma | 750 | 0.993 | 0.058 |
| Kidney papillary cell carcinoma | 719 | 0.983 | 0.062 |
| Kidney clear cell carcinoma | 784 | 0.986 | 0.060 |
| Uveal melanoma | 561 | 0.969 | 0.067 |
| Skin cutaneous melanoma | 752 | 0.983 | 0.068 |
| Rectum adenocarcinoma | 736 | 0.991 | 0.058 |
| Breast invasive carcinoma | 807 | 0.989 | 0.067 |

Table 8: Mean and standard deviation of overall predicted transcript abundance per gene (percentage of genes directed towards each transcript), considering genes with mean expression exceeding 5 TPMs in the GTEx dataset.

| Tissue | N genes | Mean Percentages | Std Percentages |
|---|---|---|---|
| Vagina | 798 | 0.970 | 0.081 |
| Uterus | 769 | 0.963 | 0.081 |
| Kidney cortex | 615 | 0.967 | 0.080 |
| Heart left ventricle | 513 | 0.938 | 0.096 |
| Cells ebv transformed lymphocytes | 683 | 0.958 | 0.089 |
| Fallopian tube | 800 | 0.969 | 0.079 |
| Pancreas | 524 | 0.997 | 0.081 |
| Whole blood | 445 | 0.907 | 0.102 |
| Heart atrial appendage | 615 | 0.960 | 0.087 |
| Prostate | 784 | 0.974 | 0.078 |
| Nerve tibial | 785 | 0.947 | 0.080 |
| Ovary | 736 | 0.958 | 0.085 |
| Pituitary | 745 | 0.964 | 0.082 |
| Breast mammary tissue | 784 | 0.974 | 0.075 |
| Lung | 809 | 0.982 | 0.080 |
| Testis | 793 | 0.911 | 0.110 |
| Cells transformed fibroblasts | 712 | 0.970 | 0.081 |
| Spleen | 750 | 0.981 | 0.079 |
| Stomach | 700 | 1.00 | 0.079 |
| Small intestine terminal ileum | 784 | 0.984 | 0.079 |
| Liver | 560 | 0.959 | 0.079 |
| Bladder | 790 | 0.972 | 0.071 |
| Muscle skeletal | 578 | 0.912 | 0.116 |
| Minor salivary gland | 760 | 0.988 | 0.073 |
| Adrenal gland | 691 | 0.975 | 0.076 |
| Thyroid | 796 | 0.979 | 0.074 |

Table 9: Comparison of additional score computation methods using POSTAR3 data. For each tumor type and method, the mean and standard deviation of the AUC calculated with the RBP scores to distinguish between classes 0 and 1 of POSTAR3 are presented.

| **Tumor type** | **Control-Kd** | | **Kup-Control** | | **Kup-Kd** | |
|---|---|---|---|---|---|---|
| | **Mean** | **Std** | **Mean** | **Std** | **Mean** | **Std** |
| Liver Hep carcinoma | 0.6987 | 0.0425 | 0.7052 | 0.0367 | 0.7058 | 0.0371 |
| Acute Myeloid Leukemia | 0.6796 | 0.0480 | 0.6709 | 0.0446 | 0.6780 | 0.0510 |
| Kidney Chromophobe | 0.7000 | 0.0607 | 0.7027 | 0.0627 | 0.7059 | 0.0616 |

Table 10: Summary of the predictive performance of the model in the RBP knockdown experiments.

| **Project** | **knockdown RBP** | **Spearman's correlation** | | **MSE** | |
|---|---|---|---|---|---|
| | | **Control** | **Kd** | **Control** | **Kd** |
| PRJEB39343 | MBNL1 | 0.777 | 0.782 | 0.263 | 0.297 |
| GSE75491 | RBM47 | 0.779 | 0.778 | 0.434 | 0.468 |
| GSE77702 | FUS | 0.703 | 0.714 | 0.585 | 0.550 |
| | TAF15 | 0.703 | 0.711 | 0.585 | 0.586 |
| | TARDBP | 0.703 | 0.703 | 0.585 | 0.618 |
| GSE136366 | TARDBP | 0.817 | 0.811 | 0.418 | 0.411 |
